# Stratification of responses to tDCS intervention in a healthy paediatric population based on resting-state EEG profiles

**DOI:** 10.1101/2022.08.09.503347

**Authors:** Paulina Clara Dagnino, Claire Braboszcz, Eleni Kroupi, Maike Splittgerber, Hannah Brauer, Astrid Dempfle, Carolin Breitling-Ziegler, Alexander Prehn-Kristensen, Kerstin Krauel, Michael Siniatchkin, Vera Moliadze, Aureli Soria-Frisch

**Author notes:** these authors contributed equally to this work (Paulina Clara Dagnino, Claire Braboszcz, Eleni Kroupi, Aureli Soria-Frisch).

## Abstract

Transcranial Direct Current Stimulation (tDCS) is a non-invasive neuromodulation technique with a wide variety of applications in both the clinical and cognitive psychology domains. As increasingly acknowledged, its effectiveness is subject dependent, which may lead to timely and costly treatments with ineffective results if this variability is not taken into account. We propose the usage of electroencephalography (EEG) for the analysis and prediction of individual responses to tDCS. In this context the application of machine learning can be of enormous help.

We analysed resting-state EEG activity to identify subgroups of participants with an homogeneous electrophysiological profile and their response to different tDCS interventions. The study described herein, which focuses on healthy controls, was conducted within a clinical trial for the development of treatments based on tDCS for age-matched children diagnosed with Attention Deficit Hyperactivity Disorder (ADHD) and Autism Spectrum Disorder (ASD).

We have studied a randomized, double-blind, sham-controlled tDCS intervention in 56 healthy children and adolescents aged 10-17, applied in 2 parallel groups over 2 target regions, namely left Dorsolateral Prefrontal Cortex (lDLPFC) and right Inferior Frontal Gyrus (rIFG). Cognitive behavioural tasks were used to both activate particular brain areas during the stimulation and to assess the impact of the intervention afterwards. We have implemented an unsupervised learning approach to stratify participants based on their resting-state EEG spectral features before the tDCS application. We have then applied a correlational analysis to identify EEG profiles associated with tDCS subject response to the specific stimulation sites and the presence or not of concurrent tasks during the intervention.

In the results we found specific digital electrophysiological profiles that can be associated to a positive response, whereas subjects with other profiles respond negatively or do not respond to the intervention. Findings suggest that unsupervised machine learning procedures, when associated with proper visualization features, can be successfully used to interpret and eventually to predict responses of individuals to tDCS treatment.

## Introduction

In recent years, Non Invasive Brain Stimulation (NIBS) techniques and, in particular, transcranial Direct Current Stimulation (tDCS)^1^,^2^. have been successfully used to study brain function, treat mental disorders and enhance cognitive functions in adults^3–12^.. This therapy can be applied as an effective treatment option^13^,^14^., which is particularly interesting in the absence of alternative effective interventions. Its application in paediatric populations however has been limited due to initial concerns^15^,^16^.. In children, stimulation parameters cannot be directly translated from studies in adult populations since electric fields generated in the brain during stimulation are affected by anatomical differences such as brain volume, skull thickness, white matter - gray matter ratio and cerebrospinal fluid (CSF) volume^17–22^.

Attention-Deficit Hyperactivity Disorder (ADHD) is one of the most prevalent neurodevelopmental chronic conditions which affects 5.9% of youth and 2.5% of adults (South America, North America, Europe, Australia, Asia, the Middle East and Scandinavia)^23^. Symptoms include hyperactivity, impulsivity, diminished attention and concentration, and vary from patient to patient. These symptoms affect in a negative way the children’s social and academic aspects of their lives^24^,^25^. Electroencephalography (EEG) has been used to characterize the ADHD population. The use of EEG-based markers overcomes some limitations of classical subjective assessments based on interviews and physical examinations. Existing reports on EEG biomarkers mention increased theta band power and decreased beta band power in the frontocentral brain region^26–28^., phase-amplitude coupling (beta-gamma) deficits in the frontal-left hemisphere^29^, and modified peak alpha frequency^30^,^31^. among others^32^,^33^.

Given the concerns that drug-based treatments of ADHD have raised in the past, there is an ongoing search for treatments with reduced or no side-effects^34^. Current drug based treatments and behavioural therapy in ADHD are also challenged by the different phenotypes and related symptoms of the pathology, as well as the complexity associated to the developing brain^35^. TDCS intervention for ADHD constitutes an interesting alternative to drug treatments^36^. However, an important factor to take into account in the development of efficient tDCS interventions is the inter-subject variability of the tDCS effect^37^,^38^. Here machine learning techniques can be of help. Past attempts have implemented unsupervised methods to characterize responses to different treatments based on clustering of genetic^39–42^, clinical^43^,^44^ and functional Magnetic Resonance Imaging^45^ data. These works report patient subgroups within a priori uniform group of patients. Such differentially responding subgroups have been also identified based on clustering of behavioural responses specifically during tDCS interventions^37^. Previous work using EEG data report prediction of response to treatment by means of Machine Learning (ML) approaches^46–48^ and stratification of clinical populations of children and adolescents^49,50^. Interestingly, variability in response to TMS and tDCS interventions in patients suffering from depression or anxiety can be explained by individual differences in baseline EEG activity^51–54^. EEG is an attractive modality for clinical application due to its lower cost and faster application time than other neurimaging modalities. In the present study, we propose to apply unsupervised clustering method to EEG data to extract so called digital phenotypes^55^ and stratify tDCS response in a healthy paediatric population.

In the present study, a tDCS intervention in a healthy paediatric population targets 2 areas of interest, namely the left Dorsolateral Prefrontal Cortex (lDLPFC) and the right Inferior Frontal Gyrus (rIFG). TDCS is applied for each area of interest both while the participants are concurrently performing a task and without any concurrent task during the stimulation. Resting state EEG was acquired before and after the tDCS intervention. We present a methodology for healthy participants stratification based on their resting state profile of EEG activity before treatment and their behavioural responses to the tDCS intervention. Behavioural metrics correspond to Accuracy and Reaction Times of Flanker task, N-back task and Continuous Performance Task (CPT). In order to determine baseline EEG profiles, we extract spectral features, which are relevant features in paediatric EEG studies^56^ and in particular ADHD^26–28^. Relative band power is calculated - this measure is preferred over the absolute band power to normalize absolute differences among participants that may present abnormalities in their anatomy^57^. To stratify participants we use the Clustering Algorithms Spectral Clustering that has good results for non-convex data^58^,59, and Fuzzy Clustering which is of interest since it can be applied to data in which boundaries are ambiguous and contain outliers^60^,^60–63^. A correlational analysis is then applied between the membership of participants to each of the a-priori electrophysiologically homogeneous groups and the a-posteriori behavioural results. Significant correlations of a particular cluster group is tested against the rest of the groups in order to confirm responders. The methodology and results we describe in this article contribute to the understanding of the variability of tDCS treatment response, with a particular focus on the specific scenarios in which the treatment is effective. We support the relationship between the tDCS response and the EEG profile on the fact that both depend on bioelectrical characteristics of the brain. Hence EEG features act as a proxy of these biophysical characteristics.

Interestingly, positive responder groups correspond to participants with a non-typical brain profile, presenting either a slowing of the EEG or a spread of increased alpha rhythm over frontal areas. This methodology can be applied later to stratify patient treatment (specifically, tDCS) enabling to focus on the specific digital phenotypes in which treatment is more probable to become effective. Personalization of treatment, applying tDCS only to groups of patients predicted to have positive results, would save costs, time and engagement to the treatment rates.

## Materials and methods

### Participants

Data come from a healthy population taking part in a clinical trial for the development of treatments for Attention Deficit Hyperactivity Disorder (ADHD) and Autism Spectrum Disorder (ASD) based on tDCS under the EU research project STIPED (Stimulation in paediatrics; grant agreement No 731827, www.stiped.eu). The study was approved by the local ethics committee of the Medical Faculty at Kiel University, Germany, and was carried out in accordance with the latest revision of the Declaration of Helsinki (Trial DRKS00008207).

A total of N=56 children aged 10 to 17 years (32 females, mean age: 14.09 years, SD: 2.1) were recruited in the Medical Faculty at Kiel University. Informed consent from the participants and their parents was obtained before the experiment. Exclusion criteria consisted of substance or medication consumption, implants and devices in the body, pregnancy, birth before pregnancy week 37, birth weight lower than 2500 grams, IQ score lower than 80 assessed with CFT 20-R Test (Culture Fair Intelligence Test)^64^, neurological and psychiatric disorders (past or present), brain surgery, presence of epilepsy in the participant or family in the present or in the past, social and health impairments as assessed with CBL (Child Behavior Checklist)^65^, FBB ADHS (Fremdbeurteilungsbogen für Aufmerksamkeitsdefizit Hyperaktivitatsstorungen)^66^ and SRS (Social Responsiveness Scale)^67^. All participants’ characteristics can be found in Tables 2 and 3 in Supplementary Information.

### Experimental Protocol

#### Experimental Design

A randomized, sham-controlled, double-blind, crossover study design was implemented. Participants came for 6 experimental sessions (see Figure 1). The first 2 sessions (T1-T2) correspond to participants screening and an optional fMRI scan. Participants then came back 4 times to receive the tDCS interventions (T3-T6) which were of 4 different types, administered in a randomized order: active tDCS with a concurrent task, active tDCS with a non-concurrent task, sham tDCS with a concurrent task and sham tDCS with a non-concurrent task. There was a minimum washout period of 7 days in-between sessions. At the start of each stimulation session, participants were asked to complete an assessment of their current mood, motivation, and any adverse event since the last session. Following the stimulation session, participants completed a questionnaire on the safety, tolerability and blinding of the stimulation.

**Figure 1.**
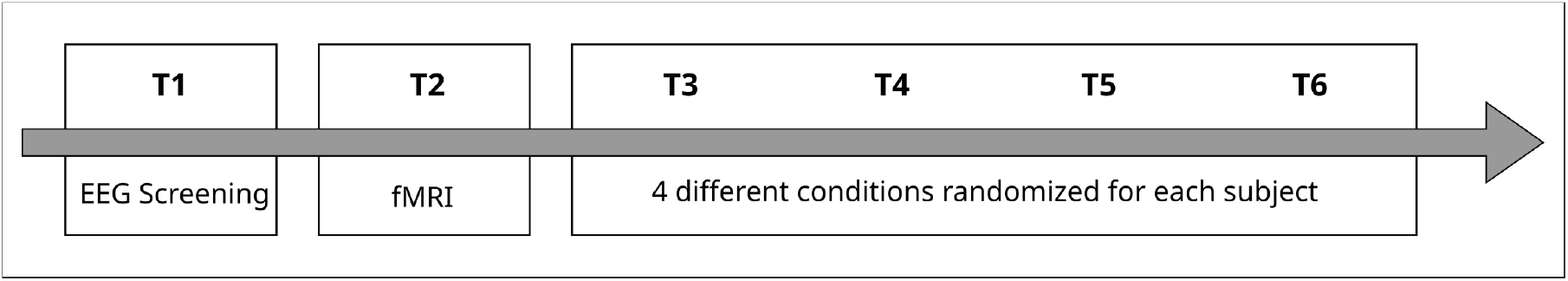
Experimental Design. Each participant is assessed at 6 time points, T1 corresponds to an EEG screening, T2 to the optional fMRI, and T3-T6 to the randomized interventions (offline/online, tDCS/sham).

Participants were randomly divided into 2 groups: participants in Group A were stimulated in the left Dorsolateral Prefrontal Cortex (lDLPFC) and performed the N-Back Task in the concurrent task condition, whereas participants in Group B were stimulated in the right Inferior Frontal Gyrus (rIFG) and performed the Flanker task in the concurrent task condition.

Each tDCS stimulation session unfolded as follows: once participants were equipped with the stimulation headset, resting state EEG was recorded during 2 minutes eyes open (EO) and 2 minutes eyes closed (EC), followed by 20 minutes of tDCS stimulation. In the during stimulation concurrent task condition, the task started after 2.5 minutes of tDCS and ended 2.5 minutes before the end of the stimulation. In the non-concurrent task condition, participants were instructed to sit and relax with eyes opened. After stimulation, a second resting state EEG was recorded (2 minutes EO, 2 minutes EC). Finally all participants performed the following cognitive tasks whilst their EEG was still being recorded: Flanker Task, N-Back Task and Continuous Performance Task (Figure 2).

**Figure 2.**
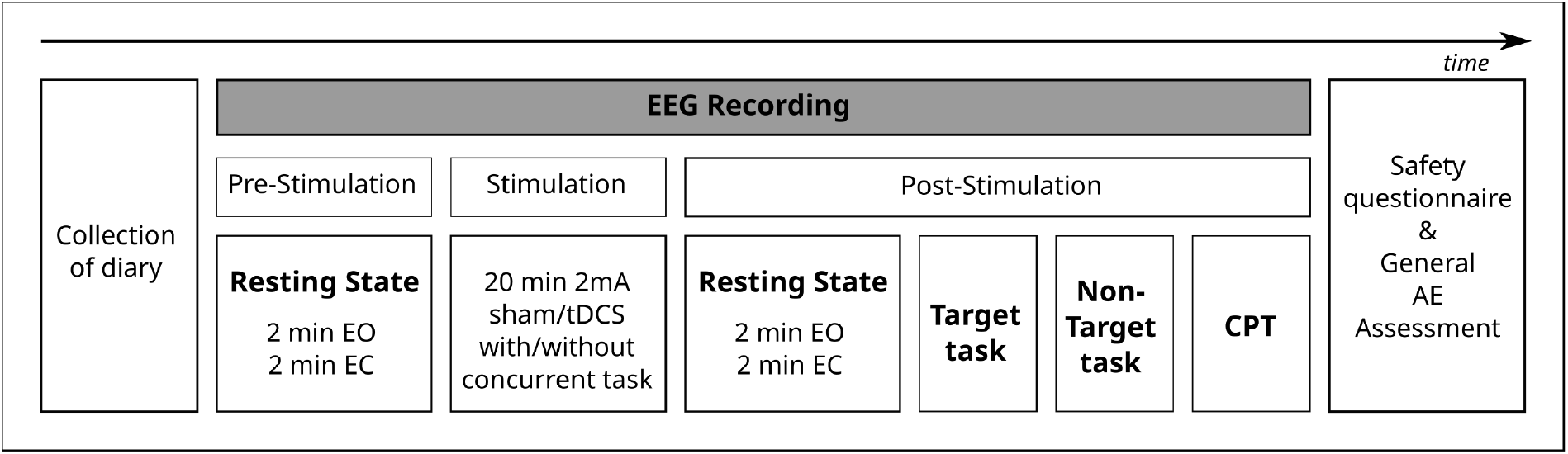
T3 to T6 Trial Design. During each trial, 2 minutes eyes open and 2 minutes eyes closed resting state EEG are being recorded before and after the tDCS stimulation. After the intervention, participants perform each of the 3 behavioural tasks whilst their EEG is still being recorded.

#### Transcranial Direct Current Stimulation

Transcranial Direct Current Stimulation (tDCS) was applied using a Starstim 32 multi-channel stimulation device (Neuroelectrics - Barcelona, Spain). The montage consisted of 5 circular electrodes (3.14 cm²) filled with gel. The total stimulation current consisted of anodal 2mA (milliAmper) during 20 minutes, with a 30 seconds of ramp up and down at the beginning and end of the stimulation (see Figure 3). The interested reader can find details on the DLPFC^68^ and the IFG^69^ montages in the referred works.

**Figure 3.**
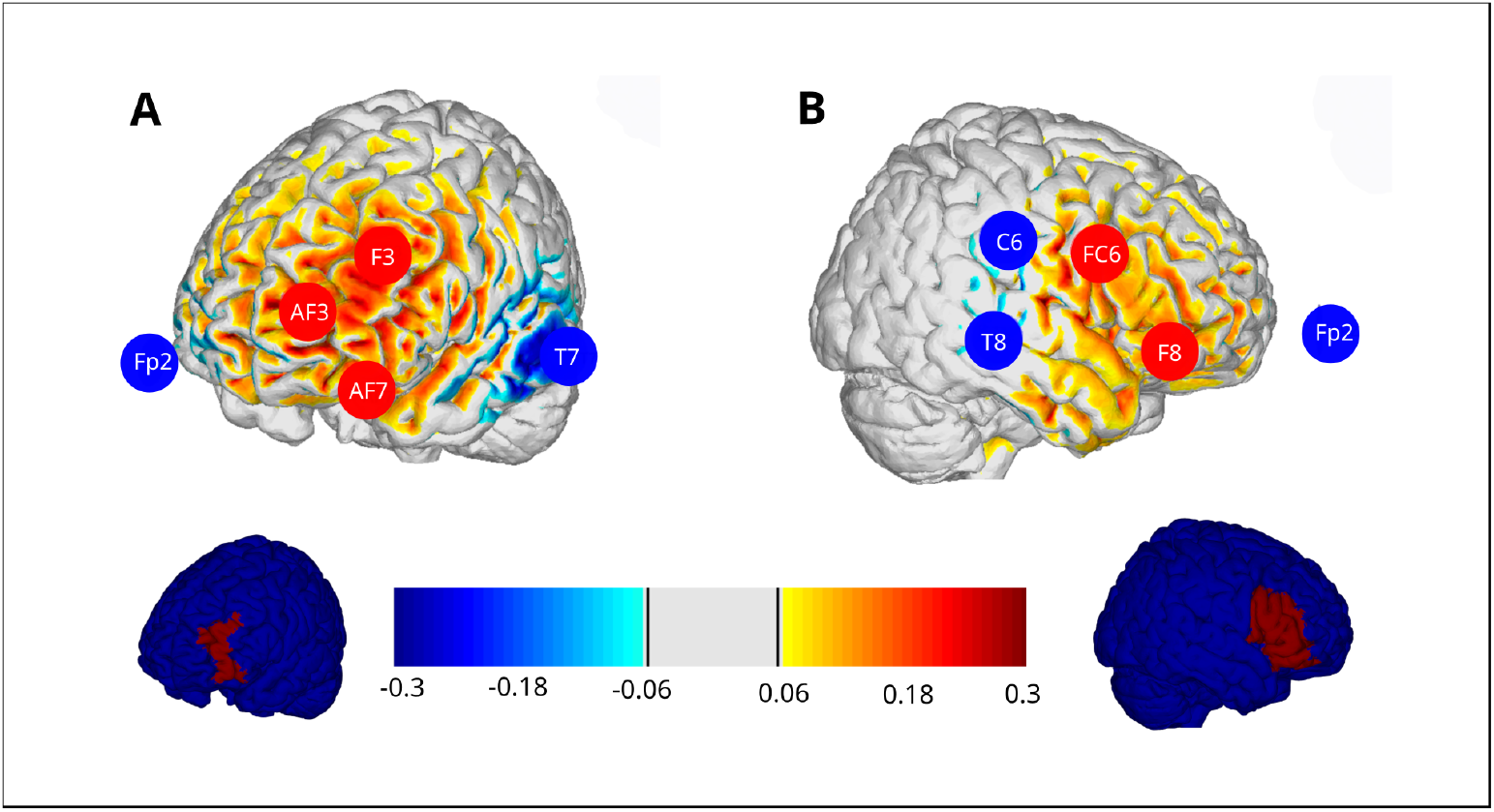
Stimulation Areas. Brain renders showing the component of the E-field normal to the cortical surface (En), positive/negative values indicate the into/out-of the cortical surface direction of the En, leading to an increase/decrease of excitability in cortical pyramidal cells. Group A of participants received stimulation in (A) left Dorsolateral Prefrontal cortex. Group B of participants received stimulation in (B) right Inferior frontal gyrus, the montage was optimized defining a target of Brodmann-areas 44, 45 and 47 on the right hemisphere.

#### EEG Recording

EEG Data was recorded using the same device used for the stimulation at a sampling rate of 500 Hz with 32 channels, following the 10-20 system: P8, T8, AF7, AF8, F8, F4, C4, P4, FC5, AF4, Fp2, Fp1, AF3, Fz, C6, Cz, C5, PO3, O1, Oz, O2, PO4, Pz, Fpz, FC6, P3, C3, F3, F7, FCz, T7, P7.

#### Behavioural Tasks

Behavioural tasks correspond to the Flanker Task, N-Back Task and Continuous Performance Test (Figure 4). These were programmed using Presentation® Software (Version 20.0, Neurobehavioural Systems, Inc., Berkeley, CA, www.neurobs.com). Behavioural performance metrics were computed in R (version 3.6.1, R Core Team^70^). The primary endpoints are the N-Back Accuracy for participants in Group A and the Flanker Accuracy for participants in Group B.

**Figure 4.**
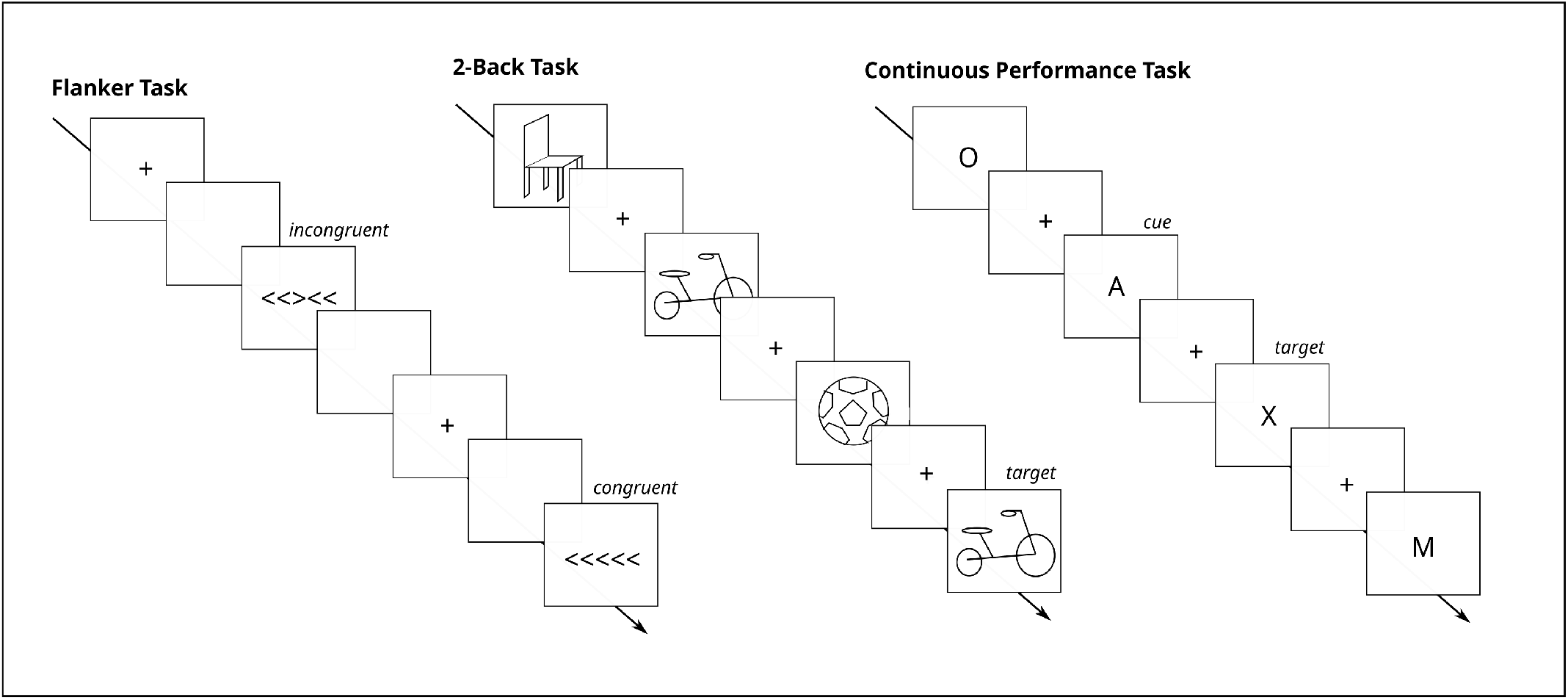
Behavioural Tasks. Examples for Flanker Task (left), 2-Back Task (middle) and Continuous Performance Task (right).

**Figure 5.**
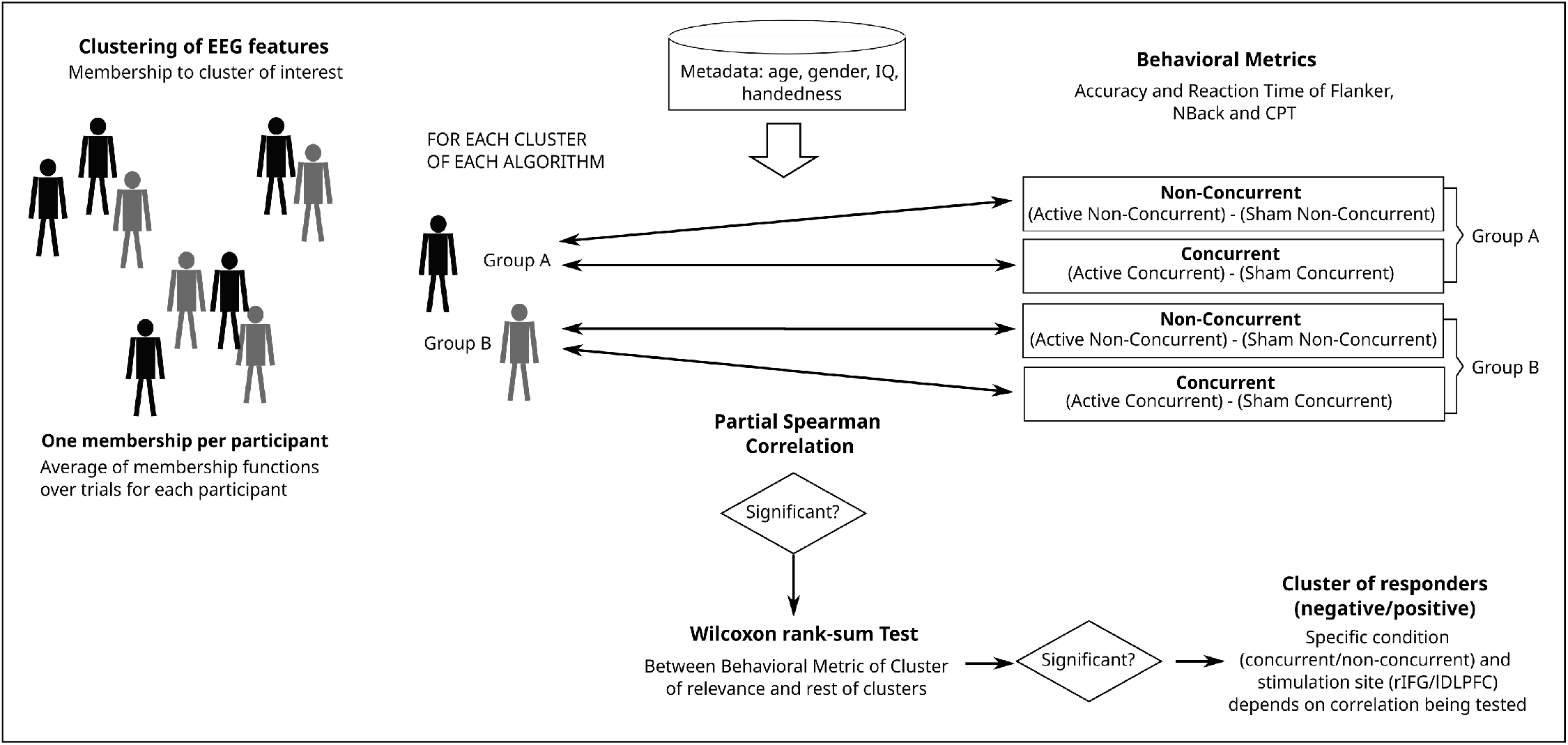
Schematic of Clustering and Wilcoxon Test Pipeline. EEG features are clustered and membership values are correlated with different behavioural metrics. If significance is reached (correlation p-value is less than 0.05), a Wilcoxon Test is done between the behavioural metric of that cluster and the rest of the points.

### Flanker Task

In the Flanker Task^71^, 5 arrows in a row were shown in which the middle one (target) points to the same or opposite direction as the rest (distractors). Participants had to press a button indicating if the target pointed left or right, with 50% of congruent trials (distractor arrows pointing in same direction as the target one) and 50% of incongruent trials (distractor arrows pointing in opposite direction as the target one). A total of 538 trials were presented in 3 different blocks with a total duration of 16 minutes. Stimuli appeared during 60 ms with an inter stimulus interval of 1676 ms. The total trial duration was of 1736 ms. Accuracy is obtained from the proportion of all hits from the total number of trials, and together with RT (in ms) used herein they represent the congruent and incongruent trials altogether.

### N-Back Task

During the N-Back Task^72^ the goal is to detect if a presented picture is the same as a picture shown before (in this case it is a 2-Back Task therefore 2 steps back). The left mouse button is pressed for a target trial, the right mouse button is pressed for a non-target trial. The task consisted of 366 trials (30% target) and lasted 16 minutes. Pictures were obtained from Stark Lab Mnemonic Similarity Task^73^ and presented during 500 ms, a fixation cross appeared during 1600 and 2000 ms between each stimulus. The total trial duration was between 2100 and 2500 ms. The performance accuracy is calculated by subtracting the errors from the number of trials, divided by the total number of trials, as a percentage. The reaction time corresponds to the time in ms to press the button in response to target hits.

### Continuous Performance Test

In the Continuous Performance Task^74^, participants were presented with a series of letters. They were shown pairs of letters in which a cue is followed by a target. They had to respond when the target appears, only when it is preceded by the cue (which is only one of the 4 existent possibilities). The cue appeared 25% of the time and the target 12,5%. Each letter was shown during 200 ms, with an inter-stimulus interval of 2200 ms. The total trial duration was of 2400 ms. The CPT accuracy corresponds to the proportion of all hits in the total number of trials, and the reaction time is the time in ms to respond in all trials.

### Signal Processing and Feature Extraction

The EEG data analysed here corresponds to the 2-minutes resting state EEG in eyes closed condition recorded before the start of tDCS stimulation in each T3 to T6 trial for each participant. Data was first band pass filtered using a 1000 coefficients finite impulse response filter with cut-off frequencies set at 2Hz and 45 Hz, demeaned and detrended. Frontal channels AF4, Fp2, Fp1 and AF3 were removed from further processing and analysis due to the presence of numerous eye movement related artifacts. High amplitude artifacts were removed using custom automated methods with 100 *µ*V as a cut-off threshold. Finally, the data was re-referenced to electrode Cz, which was then excluded from further processing. The relative band power was computed in 4-second time-windows epochs with 50% overlap in bands Delta (2-4 Hz), Theta (4-8 Hz), Alpha (8-13 Hz), Beta (13-30 Hz) and Gamma (30-45 Hz). The final relative band power values for each subject corresponded to the median of the epochs. The EEG data pre-processing and analysis was done using custom scripts in Python 3.9.

### Overview of the analytical approach

The Figure below gives a schematic of the methods used in the core analytical approach of our work. Details of each steps are given in the following Sections.

### Unsupervised Clustering Analysis

We applied 2 different unsupervised clustering algorithms to the spectral features extracted from the EEG data: Spectral Clustering (from Python scikit-learn package^1^) and Fuzzy c-Means (from Holt Skinner 2018^2^). The definition of parameters for the clustering pipeline consisted of selecting the optimal number of clusters *k* ranging from 2 to 6, and the specific parameters for each algorithm (number of components *n* for Spectral Clustering and fuzzification parameter value *m* for Fuzzy c-Means, see a more detailed description in the following Sections). These were chosen based on several internal validity measures and the visual inspection of the feature space in 2-dimensional projection based on t-distributed stochastic neighbour embedding (t-SNE), appropriate for maintaining local interactions and robust to the presence of outliers^75^. This second criteria was chosen to complement the generally used internal validity metrics we applied here since these metrics lack robustness in large dimensional spaces, similar to the one we are dealing with (5 different Band Power for each of the 27 EEG Channels).

#### Spectral Clustering

Spectral clustering has proved to be useful when the structure of the individual clusters is highly non-convex or more generally when a measure of the centre and spread of the cluster is not a suitable description of the complete cluster^59^. The algorithm is based on 3 main steps, first a matrix representation of the graph into a similarity or adjacency matrix, then a projection of this matrix onto a lower dimensional space in order to enhance cluster properties, and finally a partitioning of points through K-means clustering. The parameter to adjust is the number of components *n* which corresponds to the number of eigenvectors to use for the spectral embedding in the second step^76,77^.

#### Fuzzy c-Means

Fuzzy c-Means (FCM) is a generalization of K-mean Clustering allowing fuzzy partitions in data with non-crisp boundaries among clusters, and robustness to the presence of outliers^62^. This is enabled by allowing non-binary memberships to clusters. It looks for partitions in a set of data-points by minimizing the cost function given by:

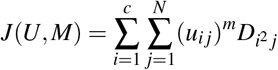

Where:

U=[u _*i j*_]_*cxN*_ is the Membership Function fuzzy partition matrix,

M=[m_1_, …, m_*c*_] is the Prototype Matrix with cluster centres,

u _*i j*_ *∈* [0,1] is the membership coefficient of the ^*th*^ object in the i^*th*^ clustering which describes the degree to which a data-point belongs to a particular cluster,

m *∈* [1,∞] is the fuzzification parameter to be explored,

D_*i j*_= D(x _*j*_, m_*i*_) is the Euclidean norm distance between x _*j*_ and m_*i*_.

#### Clustering Internal Validity Metrics

Validity measures were applied on clustering results in order to analyse algorithm performance. Generally speaking validity measures evaluate the intra- and inter-cluster distances in order to objectively quantify respectively the homogeneity and the discriminative power of the solution. The following internal validity metrics are included to evaluate the clustering result herein: Silhouette, Davies Bouldin Score (DB), Calinski-Harabasz (CH), Bayesian Information Criteria (BIC) and Inertia, implemented with Python’s Scikit-learn and SciPy packages. Silhouette is a metric which takes into account intra- and inter-cluster distances, it is the average of all data-points and its values range between -1 and 1 in which optimal highly dense clusters are given from values closer to 1^78^. Davies Bouldin Score considers the similarity between each cluster and the one it most resembles it^79^, by calculating the distance between clusters whilst considering their sizes. Values range from 0 to infinity, lower ones indicating more optimal clustering (better separation). Calinski-Harabasz calculates the ratio of sum of between-cluster dispersion and within-cluster dispersion; higher values correspond to dense and well separated clusters^77^. Bayesian Information Criterion is the likelihood of a model taking into account the variance of the data samples; the lower the value the better^80^. Inertia (or Within-Cluster Sums of Squares) is the sum of squared errors towards near centre and ranges from 0 to infinity, where the smaller is the better^81^. In summary, good clustering is represented by low values of DB, inertia and BIC, and high values of Silhouette and CH.

#### Clustering Validation and digital phenotypes

The robustness of the clustering methodology was tested through a k-hold-out workflow using 80% for parameter search. The chosen algorithms with their corresponding parameters (fuzzification parameter *m* of 1.7 for Fuzzy c-Means and number of components *n* equal to number of centres for Spectral Clustering) and number of centres *k* were applied to a subset of the data-points and then tested in the remaining points, checking whether the clusters assigned to the test set generalize to the ones obtained on the training set. This was done by first clustering the train data samples and plotting them in the feature space. The remaining test data samples labels were then assigned according to their nearest distance to the final centre previously obtained. The overlap in the feature space was studied by visual inspection and internal validity metrics were also analysed to obtain a quantitative evaluation of the generalization capability of the overall methodology.

#### Cluster Membership definition

For each participants we used 4 datasets (corresponding to each pre-stimulation EEG resting state recording in sessions T3 to T6). We thus obtained 4 different membership functions for each subject, which may present small variability due to the differences in recording time. As we are trying to establish a participant profile, we averaged the cluster membership functions of each subject over the 4 sessions. This average membership function is expected to be more robust than the individual session measures and to get rid of the temporal variability, which is not specific to a subject.

### Characterizing digital phenotypes through Statistical Analysis

#### Correlation Analysis

After clustering of the participants based on their EEG data prior to tDCS stimulation in each trial, we correlated the membership function and the behavioural data of each participant using a Partial Spearman Correlation. This type of correlation allows to take into account spurious factors that may affect the clustering result. Hence the correlation was corrected for age, sex, IQ measured by the CFT 20-R^64^ and handedness measured with the Edinburgh Handedness Inventory (EHI)^82^. A correction to take into account the multiple comparisons was applied in order to avoid random significant values. Hence the False Discovery Rate (FDR) was corrected with one of the most common methods, Benjamin and Hochberg step-up procedure (BH)^83^. The purpose of this step is to label clusters, whose members are exclusively grouped upon their homogeneity, in terms of their members’ pre-tDCS EEG resting state characteristics and group behavioural response. The behavioural response corresponds to the Accuracy and Reaction Times from the Flanker task, N-Back and CPT tasks performed after the tDCS stimulation, at the end of the trials. For each task and behavioural metric (Accuracy and Reaction Time), we computed the difference performance Delta= (Active tDCS) – (Sham tDCS). Performance Delta was computed independently for trials with a concurrent task during tDCS and trials without a concurrent task.

Since 4 experimental conditions exist (Groups A and B, concurrent/non-concurrent), for each Clustering Algorithm 4 correlations were computed between the performance Delta of each condition and the average membership to the cluster of each subject. The motivation for computing these 4 correlations is to identify the concrete protocols, i.e. stimulation site A/B and task concurrence, in which significant results are found and predict which one(s) should be used in the future. The objective is to study, for each protocol (concurrent/non-concurrent task during the stimulation), if the active tDCS stimulation improves the behavioural performance with respect to performance achieved with the sham stimulation. If there is a significant correlation (results are considered statistically significant if the correlation p-value is less than 0.05), we can establish a relationship between the EEG profile and the actual difference of behavioural results between the active and sham stimulation. We denote this difference as the Delta in performance. If this were not the case, then we would be looking for significant correlations including both active and sham conditions without any distinction among them. Positive responders are detected for positive Accuracy correlations and negative RT correlations, since for the former it means that the active condition has a higher accuracy than the sham condition, whereas the opposite is represented in the latter for RT. On the other hand negative responder clusters are defined by presenting a negative correlation with Accuracy measures and positive correlation with RT ones.

#### Wilcoxon Test

If any of the 4 computations showed a significant correlation (positive or negative), then we analysed the difference in behavioural response for the members versus the non-members of the cluster. To do so, a rank-sum Wilcoxon Test was calculated for the behavioural metrics, between each cluster samples and the samples belonging to the remaining clusters. Benjamin and Hochberg step-up procedure (BH)^83^ was used to correct for multiple comparisons. In this part of the analysis we want to investigate if there are significant differences between the behavioural outcomes for each specific cluster, once the cluster has shown a correlation between the membership to that cluster and the corresponding behavioural metric. Hence, for each test in a cluster with a significant correlation, the significance of the difference in behavioural Delta of that subject cluster members was evaluated with respect to the subjects belonging to the rest of the clusters. We illustrate this operation with an example: If a significant correlation is found for Cluster 1 in behavioural metric Accuracy Flanker – considering the membership values of all data-points to that specific Cluster 1 – then a Wilcoxon test is performed on the Flanker Accuracy Delta between the subjects that belong to Cluster 1 and the rest of the subjects).

#### Membership to Clusters

The membership of each data samples to each of the cluster groups were used for the correlation. This was motivated to avoid the binarization of the data. By working in a real-valued domain for the membership values we are able not to treat equally points which are near to the cluster centres and the points that are near the boundaries. This is expected to increase the robustness of the analysis pipeline, given the fact that early binarization thresholds largely influence final results.

For the Wilcoxon test, since we were calculating the significant difference between the subject samples in one cluster and the rest of the subject samples, we needed to determine to which final cluster a subject belongs to. This was done through the argmax function over its membership functions, i.e. determining the cluster with a highest membership value (e.g., if point X has a membership value of 0.2, 0.3, 0.4 and 0.1 for Clusters 1, 2, 3 and 4 respectively, then it would belong to Cluster 3).

#### Validation of the Statistical Analysis

The robustness of the statistical analysis procedures, i.e. Correlation and Wilcoxon Test, was validated through repeated hold-out. The correlation was computed 1000 times with different percentages of participants (80%, 60% and 40% of participants). The average and standard deviation of the strength of the correlation and its p-value were computed over the repetitions. Analogously the average and standard deviation of the p-value resulting from the Wilcoxon test was computed. For the correlation we have conducted an additional validation step in which the error between the linear regression originally obtained with 100% of subjects and the samples of the subjects left out in the validation was calculated.

## Results

### Feature Space Definition

We computed the Relative Band Power in bands Delta, Theta, Alpha, Beta and Gamma for each electrode resulting in a feature space of 135 components. After removal of artifactual signals and due to the absence of some trials or due to dropout subjects, 206 data-points were used. Figure 6 shows the optimal feature space visualization with t-SNE in 2 dimensions, given by a perplexity index value of 30^75^. It can be seen that the neurophysiological profiles of all trials for all participants are very close to each other, meaning that the intra-subject variability is small enough to consider robust the clustering results.

**Figure 6.**
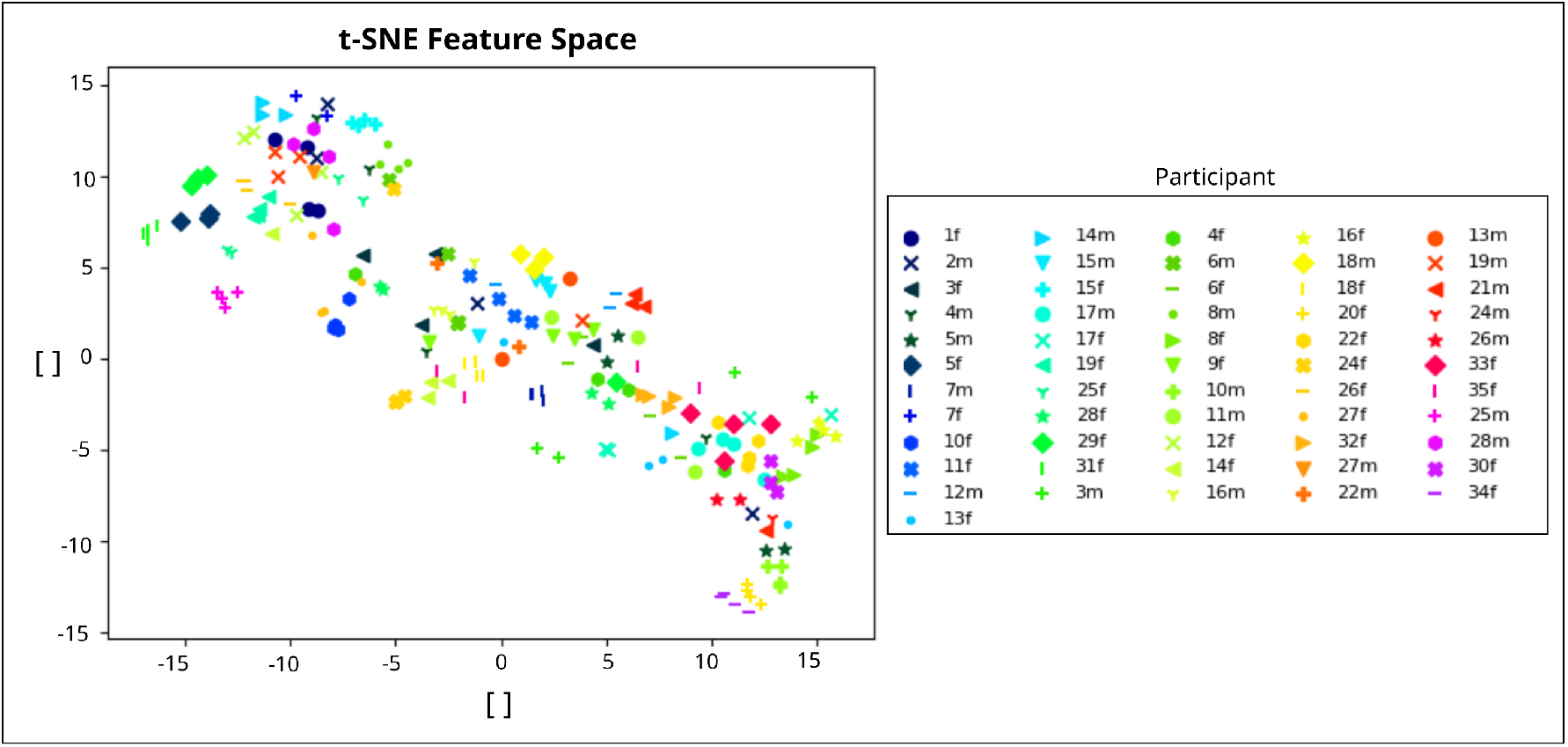
Feature Space. Visualization of EEG data-points with t-SNE in 2 dimensions. Each point corresponds to a specific trial for each participant, the features of each data-point are the RBP in each channel.

### Clustering Results

We implemented the different clustering algorithms to establish homogeneous subgroups among subjects based on their resting-state EEG spectral features before intervention. The evolution of the validity measures over the different parameters do not offer a clear criterion for parameter selection. We believe this may be due to the high dimensionality of the feature space, i.e. the “curse of dimensionality”^84^, which is not clearly tackled by the validity measures used. In spite of this, we observe some singular points in the evolution of the selected validity measures over clustering parameters (see Figure 7 B). Because of this lack of a clear objective criteria for parameter selection exclusively based on the validity analysis, we have taken into account an additional qualitative evaluation criteria based on the visualization of the t-SNE projection of the feature space, showing well separated and clear cluster clouds depicted in Figure 7 A. For the Spectral Clustering, a number of components *n* equal to the number of centres was used, as it generally corresponds to the default parameter. This gives better results than a number of components *n* equal to 100 applied as an example for extreme values, and gives similar results to smaller values such as 2, 6 and 8. For FCM the chosen parameter was a fuzzification parameter *m* of 1.7 since it is the furthest away value from 1 so as to be more dissimilar to K-means, with optimal values for the internal validity metrics and visualization in the feature space.

**Figure 7.**
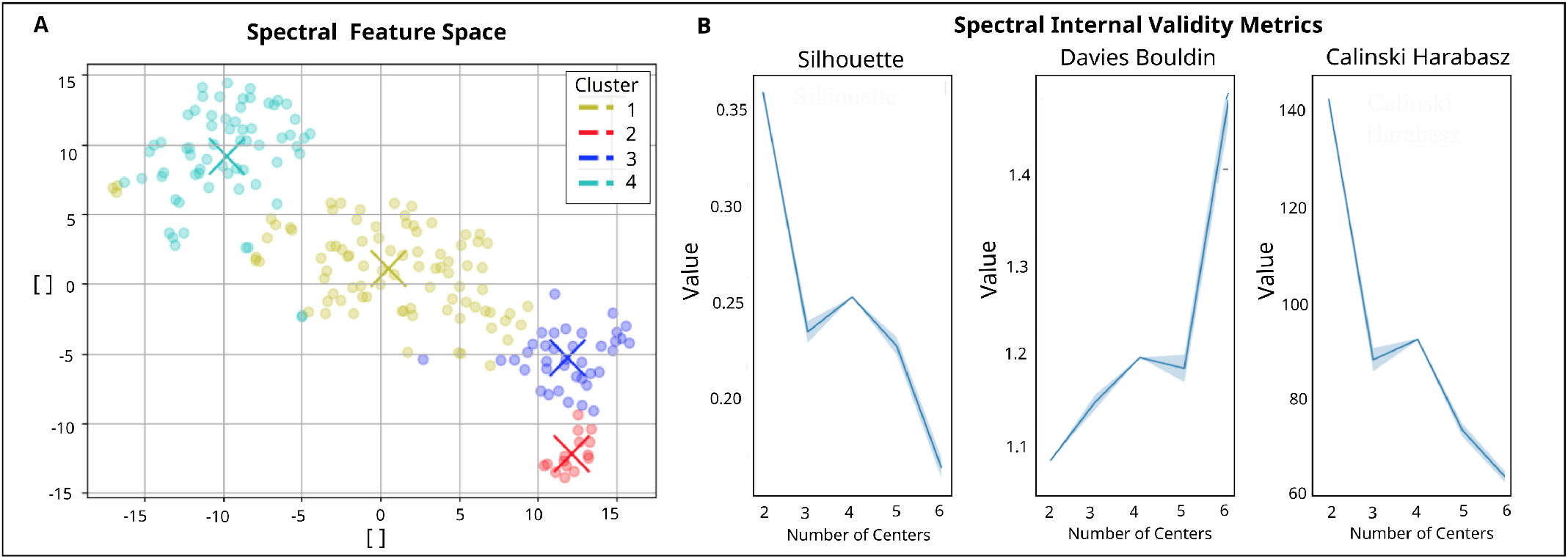
Clustering Results for Spectral Analysis. The left plot shows the feature Space, colourcode represents each cluster. From all data-points, 42,7% lie in Cluster 1, 7.3% in Cluster 2, 18.4% in Cluster 3 and 31.6% in Cluster 4. The right plots show the evolution of the internal validity metrics Silhouette and Davies Bouldin when changing the number of centres. An elbow at k=4 can be seen.

Concretely we can observe some singular points, i.e. “elbows”, at the cluster centre values of 4 for Spectral Algorithm which correspond to the optimal values. For the case of FCM Algorithm it is optimal in *k*=5. In order to have the most amount of data-points in each cluster cloud, and since it shows clearly defined clusters in the feature space result from both algorithms, we eventually defined the number of cluster centres as *k*=4. The following analysis corresponds to the results obtained for the Spectral Algorithm (FCM results for *k*=4 can be found in Supplementary Information). Since the clustering output visualized in the feature space shows similar results for both algorithms, a possible hypothesis for the differences in point cluster assignment between Spectral and FCM is the difference in the way the membership functions are computed in each algorithm.

The robustness of the clustering methodology was analysed through a k-hold-out using 80% for parameter search. The validation is successful since a visual overlap can be observed between the clusters assigned to the set not used in the parameter search (Figure 6 A in Supplementary Information). Moreover, the internal validity metrics follow the same trend in both data subsets (Figure 6 B in Supplementary Information).

### Electroencephalographic Profiles

In order to characterize each of the clusters in terms of electrophysiological profiles, the prototypes generated for each cluster of each algorithm were used. The electrophysiological profiles are set up as a combination of a representation in the frequency and spatial domains. First, the power spectrum for each cluster prototype is plotted (Figure 8) using the mean of occipital and parietal electrodes (O1, O2, Oz, P8, P4, P3, P7, PO3, PO4, Pz), where the alpha peak during eyes closed should be more characteristic. Topoplots are used for the representation of prototypes in the spatial domain (Figure 10). They are visualized calculating the mean of the Relative Band Power of all of the data-points belonging to each cluster. Since Cz was used as reference and removed from the analysis, for topoplot visualization it is interpolated and calculated as the mean of the surrounding electrodes C3, C4, Pz and FCz. As a result some differences might appear in this region.

**Figure 8.**
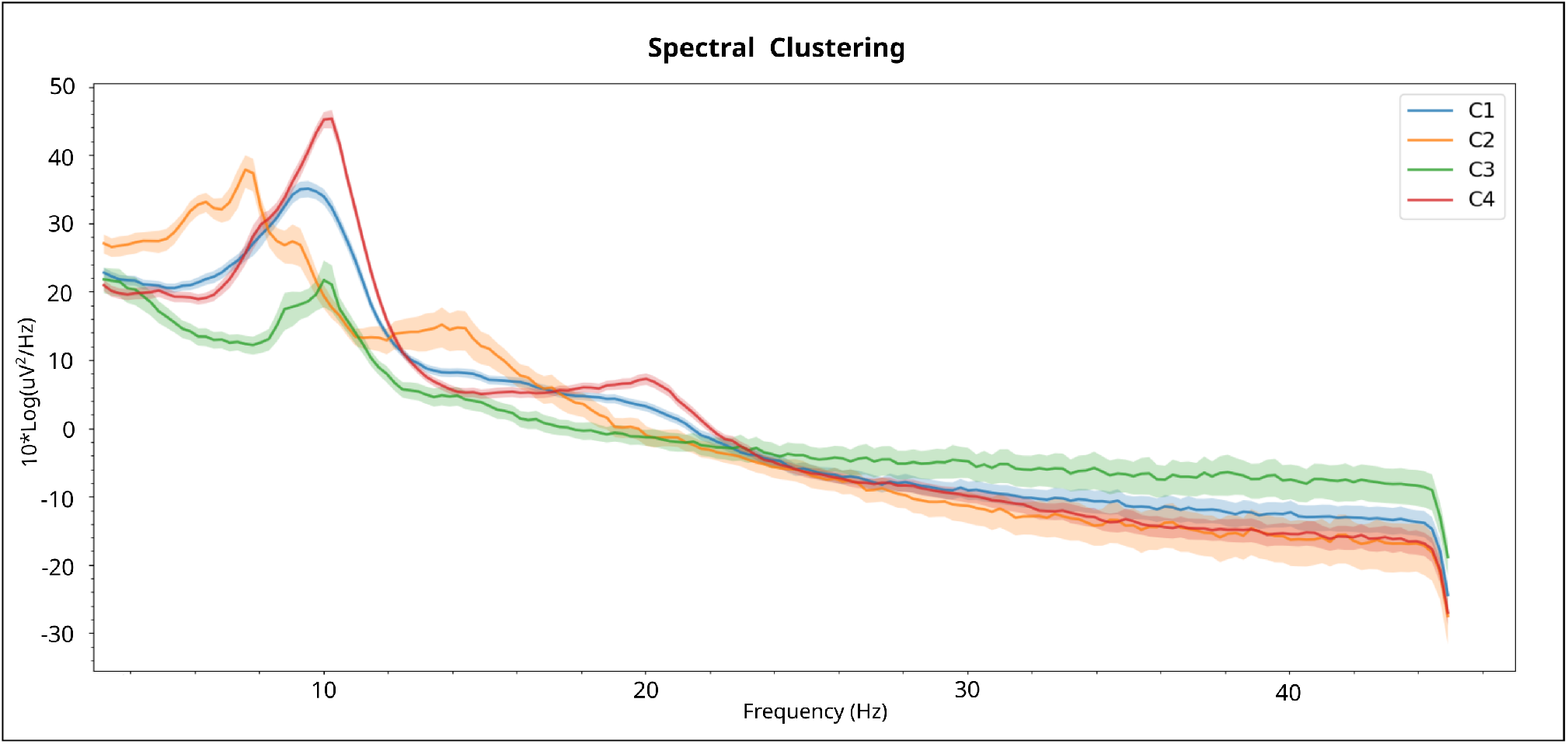
Power Spectrum Plot of each Cluster for Spectral Algorithm Results.

**Figure 9.**
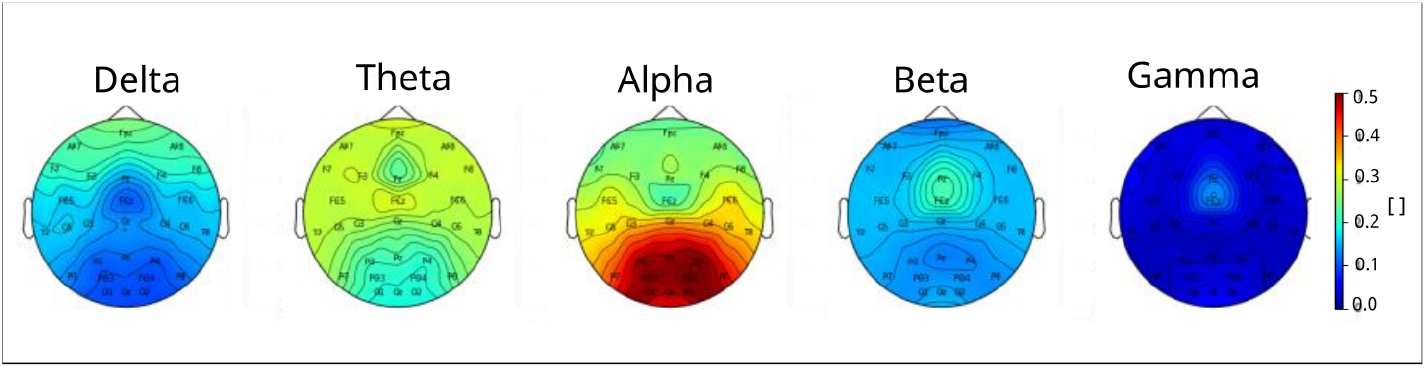
Overall Population Topoplot.

**Figure 10.**
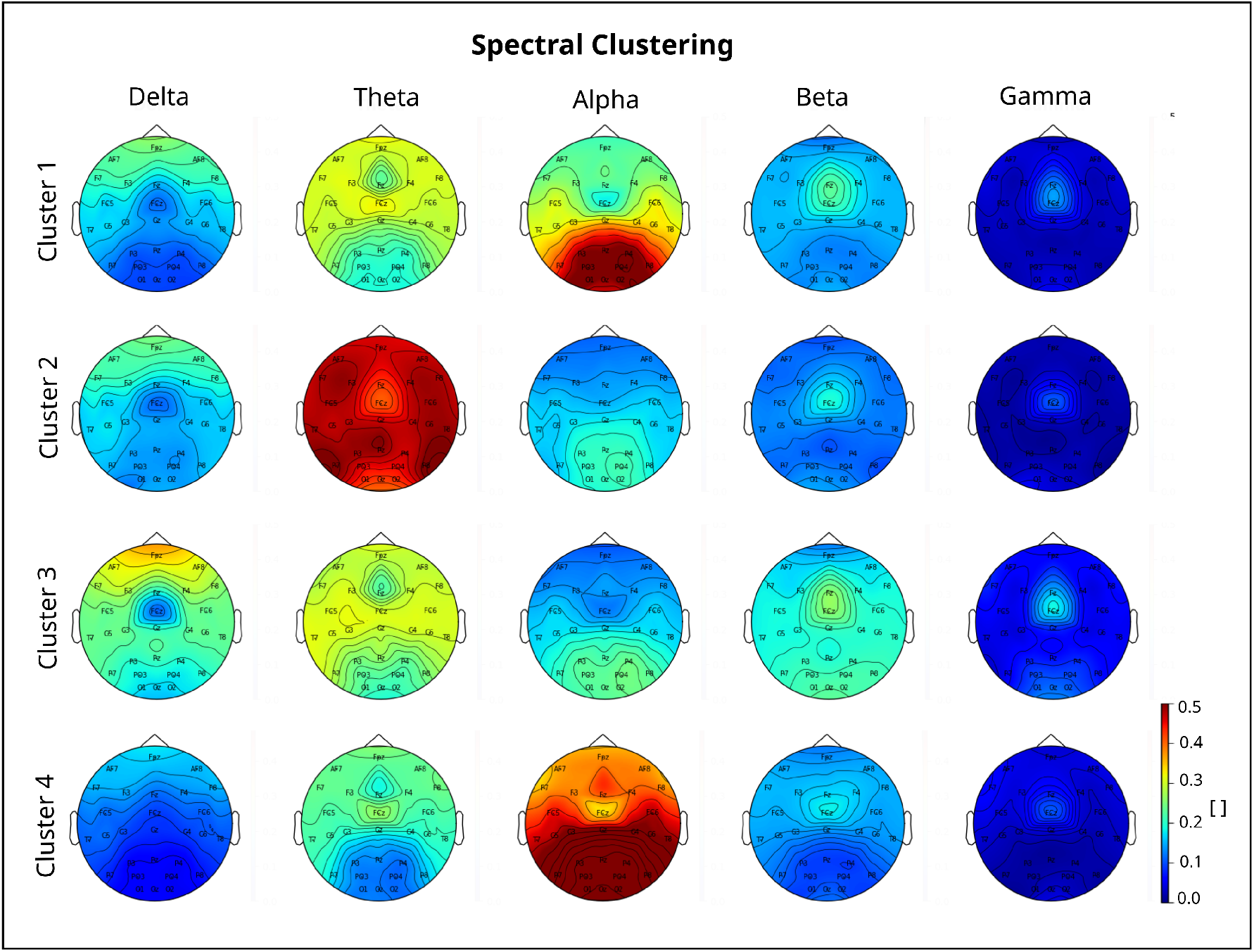
Power Spectrum Plot of each Cluster for Spectral Algorithm Results.

Cluster 1 presents a typical PSD and scalp power distribution for eyes-closed, with alpha power localized over posterior regions. Cluster 2 presents a very clear slowing in both the PSD and the topoplots, from the Alpha band towards the Theta band. Cluster 3 presents low amplitude in 10 Hz and high amplitudes at higher powers in the PSD, and a slowing in the topoplots. Cluster 4 presents the highest amplitude in 10 Hz and a peak in 20 Hz in the PSD, and the difference between frontal and parietal areas for the Alpha band in the topoplot is smaller compared to the rest of the clusters. As a remark, Cluster 1 topographical map of activation is the most similar to the ones of the overall study population (see Figure 9).

### Behavioural Correlation and Wilcoxon Test

We applied a correlation analysis between participants’ membership to each cluster and the post-tDCS behavioural results of Accuracy and RT of Flanker, N-Back and CPT tasks. Due to low variability in the baseline EEG spectral activity of each participant (Figure 6), the average of the membership function over all trials of each participant was computed in order to eliminate trial dependency, and obtain a more robust characterization with just one point per subject. As a result, the same membership values, corresponding to the average of all trials for a given subject, are used for the correlation analysis with the performance metrics in each trial of the corresponding subject. Eleven subjects were excluded due to incorrect randomization and/or dropouts during the trials, and behavioural outliers are removed for each metric using a threshold of +-2.5 standard deviations. The following matrices (Figure 11) show the strength of the correlations and the range of the p-val when significant (* - 10%, ** - 5% and *** - 1%) after correction for multiple comparison with FDR (24 corrections for Group B, 20 corrections for Group A). A specific example can be seen in Figure 5 A in Supplementary Information.

**Figure 11.**
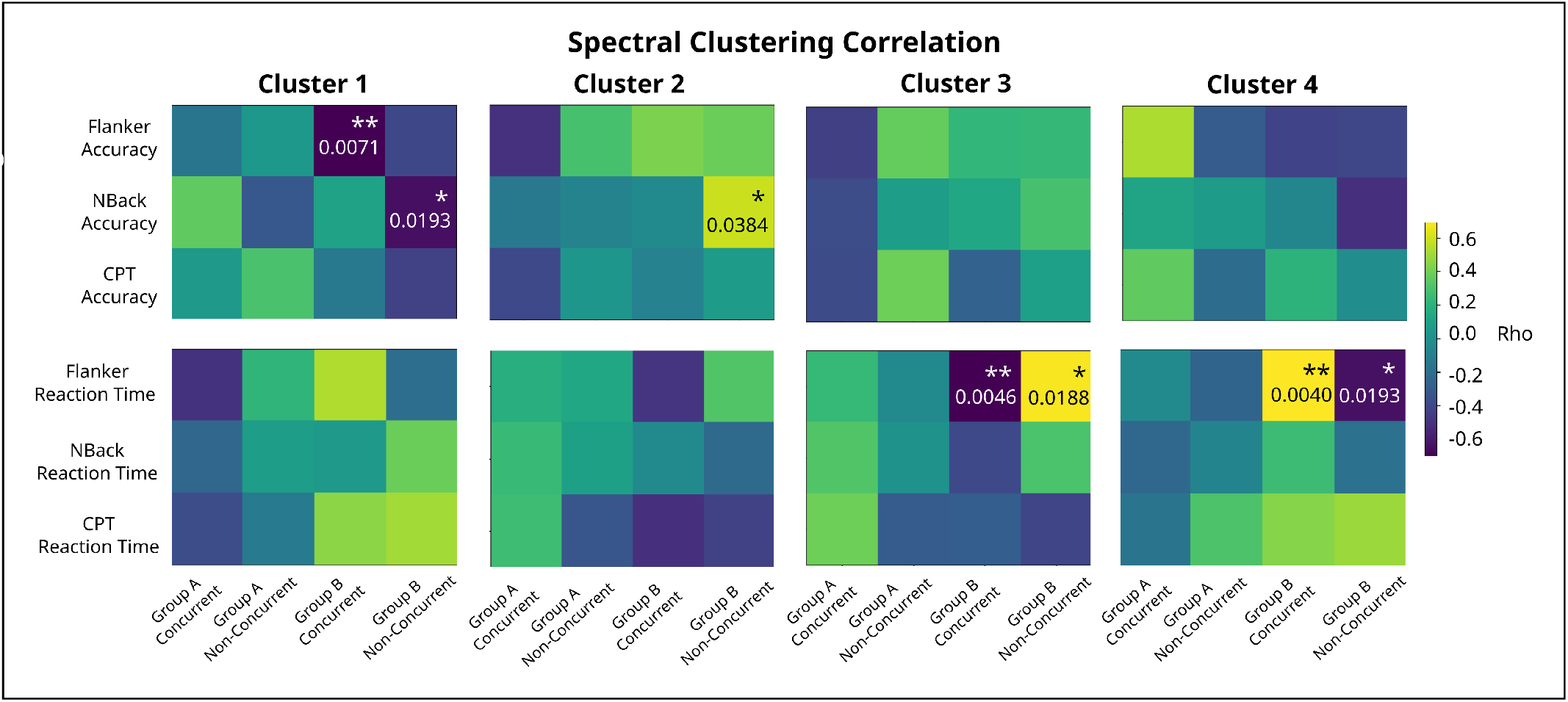
Correlation Results for Spectral Clustering. Each matrix row corresponds to a specific behavioural task endpoint, and each matrix column corresponds to a particular treatment protocol. The colourbar represents the sign of the correlation (positive or negative) and p-values are written in the significant cells. For Group B, 24 corrections are done; for Group A, 20 corrections are applied.

After computing the rank-sum Wilcoxon test only in the significant correlation results, responder clusters (positive or negative) were confirmed from the ones that also give significant differences in this test. Corrected results for multiple comparisons for the Wilcoxon analysis, as well as number of corrections, are presented in Table 1. A specific example can be seen in Figure 5 B in Supplementary Information.

**Table 1.** Spectral Significant Responders. These results correspond to significant correlations and significant Wilcoxon tests. In parenthesis are: (1) type of response (positive or negative), (2) Wilcoxon p-values. There are significant correlations only in Group B, therefore for Wilcoxon Test 4 corrections are done for Non-concurrent Group and 3 for Concurrent Group. Stimulation protocol Group B (rIFG), offline mode (non-concurrent task) presents positive response in Clusters 2 and 4 measured with N-Back Accuracy and Flanker RT respectively, and negative response in Cluster 3 assessed with Flanker RT. Stimulation protocol B (rIFG), active mode (with Flanker concurrent task) presents negative response in Clusters 1 and 4 measured with Flanker Accuracy and Flanker RT respectively, and positive response in Cluster 3 assessed with Flanker RT.

| Cluster | Group A, Concurrent | Group A, Non-concurrent | Group B, Concurrent | Group B, Non-concurrent |
| --- | --- | --- | --- | --- |
| 1 |  |  | Flanker Accuracy (-, 0.0150) |  |
| 2 |  |  |  | N-Back Accuracy (+, 0.0345) |
| 3 |  |  | Flanker RT (+, 0.0150) | Flanker RT (-, 0.0433) |
| 4 |  |  | Flanker RT (-, 0.0402) | Flanker RT (+, 0.0345) |

The robustness of the correlational analysis was tested iterating 1000 different percentages of the data set (specifically 80%, 60% and 40%). We find a trend for the coefficients and p-values of both correlations and Wilcoxon tests. Overall, the sets have same direction of correlation as the whole dataset, and the strength goes diminishing in the order 80%-60%-40%. With respect to the correlation p-value, it is almost always significant (correlation p-value is less than 0.05) for the 80% dataset. In the 60% dataset it is generally non-significant although very close to the boundary (correlation p-value near 0.05) and for the 40% it is never significant. In both cases, the standard deviation increases when the percentage of data-points decreases. These takeaways have similar trends for the analysis using Wilcoxon test. For the case of the distance to the original fit line (with the 100% of data-points), the mean is very similar in all percentages of the dataset, whereas the standard deviation increases for decreasing percentages. Overall, results show that our correlation method is robust and that the sample size used is the minimum required for obtaining these results.

Since it has been seen that subjects who better respond to treatment correspond to those who start in worse conditions^85–87^, an extra analysis was done in order to discard the differential response in terms of this factor. Hence we further studied whether there is a relationship between baseline behavioural measures (at T3) and treatment response. The following Figure 12 shows the behavioural response for the baseline (T3) of the main endpoints (Flanker Accuracy and N-Back Accuracy) as well as for the RT of those tasks. No cluster structure can be seen on the behavioural metric at baseline. Therefore we can state that the cluster structure is solely due to homogeneous electrophysiological profiles and not at differential baseline response in the tasks.

**Figure 12.**
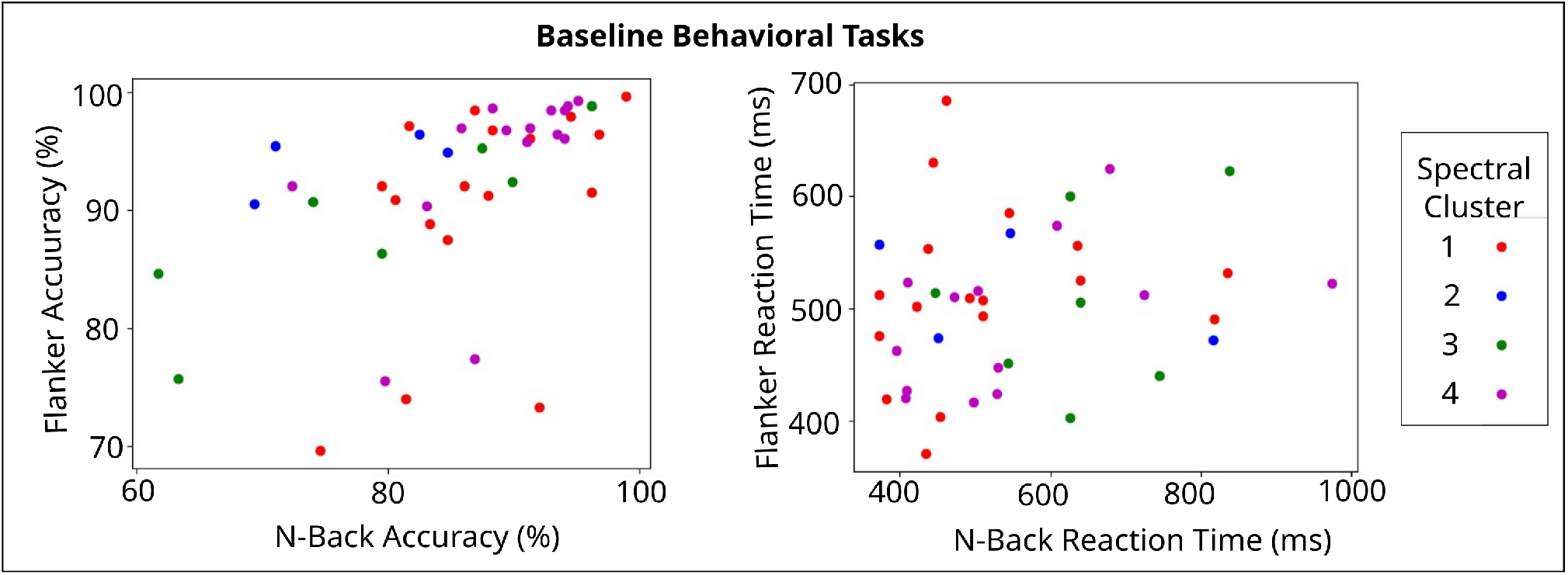
Baseline behavioural tasks distinguished by Spectral Algorithm Clustering for (A) Accuracy and (B) Reaction Time.

## Discussion

Our work provides a novel approach to study the effects of multichannel anodal tDCS treatment in healthy children and adolescents with different specific brain profiles. This is of interest given the limitations of existent treatments in paediatric clinical populations, e.g. ADHD and ASD for which new treatments with no side effects^88^ are being developed. Furthermore the prognostic value of such approaches can be exploited in the context of the development of reliable and well-defined personalized protocols. Given that significant positive response to treatment is not always guaranteed, such an approach can be of enormous value for the cost-effective development of new treatments.

By subgrouping healthy controls in those with typical versus non-typical EEG profiles, a differentiated response can be found related to the specific non-typical electrophysiological profiles. Profiles (summarized in Figure 13) are characterized only by homogeneous resting-state EEG spectral features and not by behavioural response to tasks before each intervention. This confirms our hypothesis that the electrophysiological profile can be used as a proxy of different biophysical characteristics of subject brains. The fact that these are almost identical at 4 different measurement time points as shown in the intra-subject variability analysis further supports the fact that the electrophysiological features used are subject specific and do not correspond to a difference in brain state, which is known to influence as well the response to Non Invasive Brain Stimulation (NIBS). We propose the usage of the stratified electrophysiological responses as shown herein as digital phenotypes for the characterization of tDCS response.

**Figure 13.**
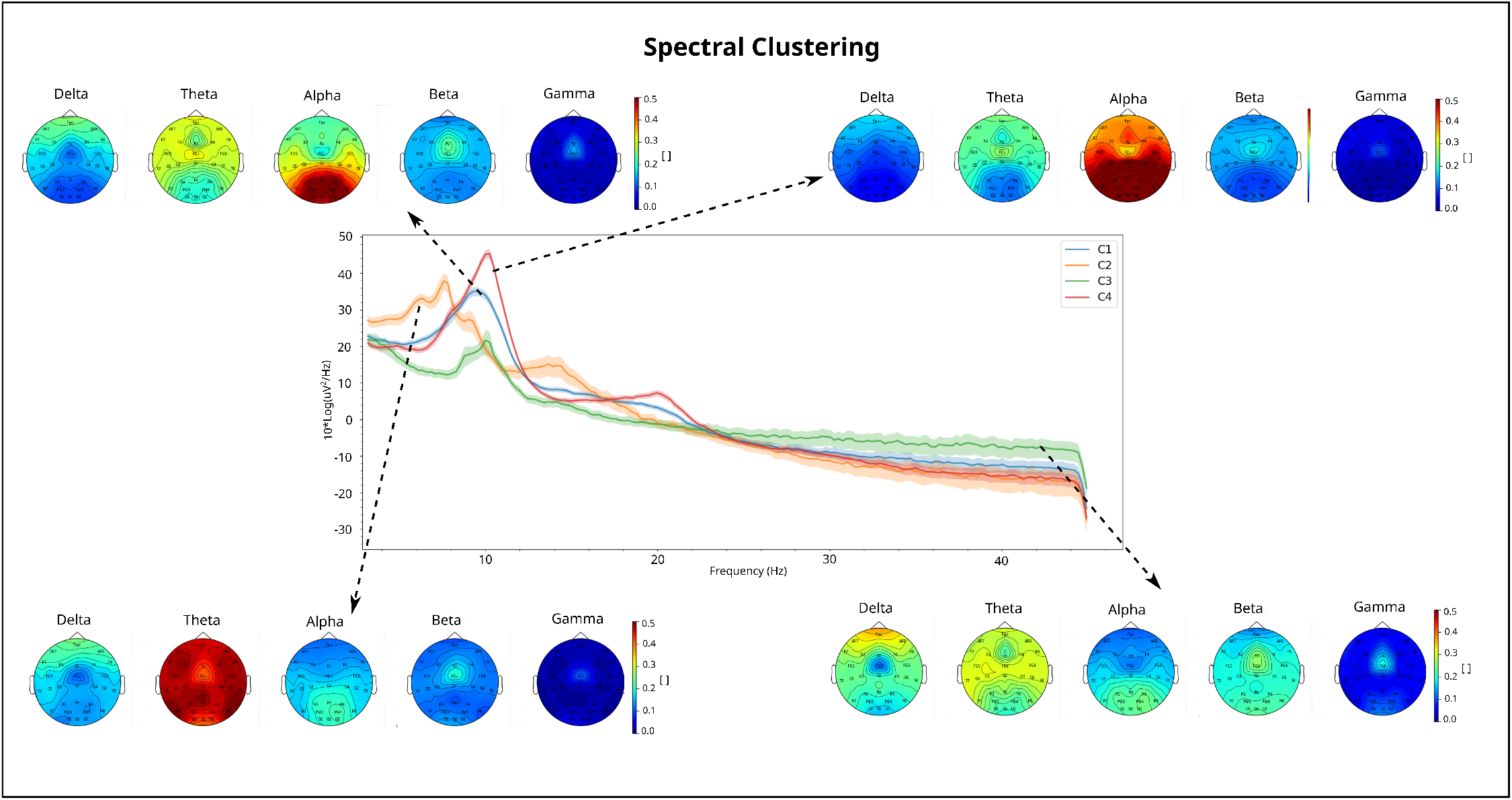
Summary of EEG Prototypes. Topoplots and PSD for Clusters that present a significant response in Spectral Algorithm. Cluster 1 has a negative response to rIFG stimulation with Flanker concurrent task, measured with Flanker Accuracy. Cluster 2 has a positive response to rIFG stimulation with no concurrent task, measured with N-Back Accuracy. Cluster 3 has a positive response to rIFG stimulation with Flanker concurrent task, measured with Flanker RT; and a negative response to rIFG stimulation with no concurrent task, measured with Flanker RT. Cluster 4 has a positive response to rIFG stimulation with no concurrent task, measured with Flanker RT; and a negative response to rIFG stimulation with Flanker concurrent task, measured with Flanker RT.

It can be observed that Group B (rIFG stimulation) is the only one with responders (positive and negative), and Group A does not have any significant correlations supported with significant Wilcoxon-test results. This suggests an advantage in stimulating the rIFG over the more traditional approach of lDLPFC stimulation even when participant stratification is applied. In almost all groups of participants, the behavioural task showing changes is the Flanker task when the applied stimulation montage targets augmenting activity of the rIFG. This result is consistent with current understanding of the role of the rIFG, associated with response inhibition and behavioural control^89^,^90^.

Future work could be done in order to establish a relationship between the electrophysiological profiles and functional outcomes. A hypothesis would be based on the transfer stimulation effects from stimulated WM network involved in the Flanker task to the WM network components involved in N-Back task^68^,^91–93^, and the response dependency to the brain’s stage of development since for example Clusters 3 and 4 have different neurophysiological characteristics and show opposite results^68^,^94^.

In summary, the effect of brain stimulation (and its parameters including location and presence/absence of concurrent task during the protocol), measured by different behavioural measures, depends on the individual digital phenotype of the subjects undergoing the stimulation intervention. The robustness of the overall developed methodology was validated in 3 independent tests. First, the feature space of all participants was visualized showing that all trials of pre-treatment EEG for each subject lie near and meaning that their baseline neurophysiological activity was similar over the different trials. Second, the k-hold-out methodology applied to the clustering pipeline demonstrates the robustness of the clustering algorithms in our specific dataset. This validation let us ensure to a certain extent the generalization capability of the clustering for generating similar digital phenotypes on unseen data. Finally, the k-hold-out methodology applied to the correlation of the clustering and wilcoxon test pipeline shows a decreasing trend with decreasing sizes of our dataset. Moreover it shows also that the behavioural response differences are robust enough to resist a reduction of 20% in the data set sample size.

Some limitations arise from the fact that the dataset is relatively small compared to the large amount of features analysed, resulting in a very high dimensional space for clustering studies and providing correlations with low clinical utility. In order to study feature importance, Random Forests^95–98^, Principal Feature Analysis^99^, permutation analysis^47^ and other feature selection algorithms^96^,^100^ could be explored to reduce the number of features, as well as other stratification methods for unsupervised clustering and machine learning techniques robust to high dimensionality^101–105^. Additionally, Clustering behavioural metrics other than RT and Accuracy or specific power bands and channels of interest in the ADHD community could be studied. Proposals are Theta Band Power which is involved in cognitive processes^106^ and for which increased values reflect the maturational delay^47^, or the Beta Band Power, particularly in frontal areas which is associated with executive control and working memory^105^,^107,108^.

## Conclusions

The advanced clustering methodology we developed shows promising results in the context of broad and heterogeneous EEG signals^109^. Moreover the use of unsupervised learning approaches allows to obtain an actual data-driven stratification, since the learning procedure is freely run without the constraints imposed by the ground truth. This is an important factor given the large misdiagnosis rates in the target diseases^110^,^111^, especially important in ADHD^111^. In case of having used a supervised approach, a false ground truth might have driven the class structure wrongly and hinder the accuracy of the stratification.

Stimulation brain target areas studied were lDLPFC and rIFG, paired or not with concurrent tasks N-Back and Flanker Tasks respectively. Effectiveness of the treatment was assessed by different behavioural measures, in particular RT and Accuracy of the aforementioned tasks, and CPT. The digital phenotypes obtained from pre-treatment EEG, as assigned by the clustering procedure described throughout the work, can be then used to predict the response of individuals to particular stimulation protocols. We have successfully developed a procedure for labelling clusters in terms of their behavioural response in different tasks. Hence a rank-sum Wilcoxon test is succesfully applied for validation and consistency of results.

The correlation analysis shows that intervention is influenced by an individual’s brain state, meaning that tDCS stimulation is state-dependent, indicating the relevance of patient screening before undergoing a treatment. Interestingly, positive responder groups correspond to participants with a non-typical brain profile, presenting either a slowing of the EEG or a spread of increased alpha rhythm over frontal areas. Although this finding is extremely interesting from a neurophysiological point of view, we focus in this communication on the novelty of the methodology of our work since a larger cohort of participants would be needed for more generalizable results and higher clinical utility.

## Acknowledgements

This study was partially supported through project STIPED. STIPED project has received funding from the European Union’s Horizon 2020 research and innovation programme under grant agreement No 731827. Author C.B work has received funding from the European Union’s Horizon 2020 research and innovation programme under Marie Skłodowska-Curie grant agreement No. 801342 (TecniospringINDUSTRY) and the Government of Catalonia’s Agency for Business Competitiveness (ACCIÓ). We give special thanks to Maike Splittgerber and Vera Moliadze for conducting the EEG recordings and data management. Professor Astrid Demple, for assistance with statistical analysis. We acknowledge the participants in this study.

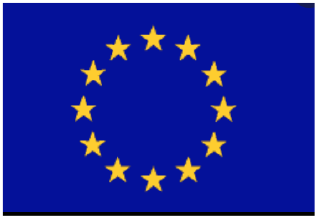

## Author contributions statement

Study concept and design: M.Sin., K.K., V.M. Conduction of experiment and Data collection: M.S., V.M. Analysis: P.D., A.S.F., E.K., C.B. Interpretation of results: P.D., A.S.F., E.K., C.B. Analysis supervision: A.S.F., E.K., C.B. Drafting of the manuscript: P.D., A.S.F., E.K., C.B. All authors reviewed the manuscript.

## Additional information

Neuroelectrics is the company that manufactures the Starstim devices used for stimulation and recording of EEG analysed in this work. Giulio Ruffini is a co-founder of Neuroelectrics. We, all authors, declare there are no conflict of interests with this publication and we have all approved this final version.

## Footnotes

1 https://scikit-learn.org/stable/

2 https://github.com/holtskinner/PossibilisticCMeans

