## Supplementary Information for "Stratification of responses to tDCS intervention in a healthy paediatric population based on resting-state EEG profiles"

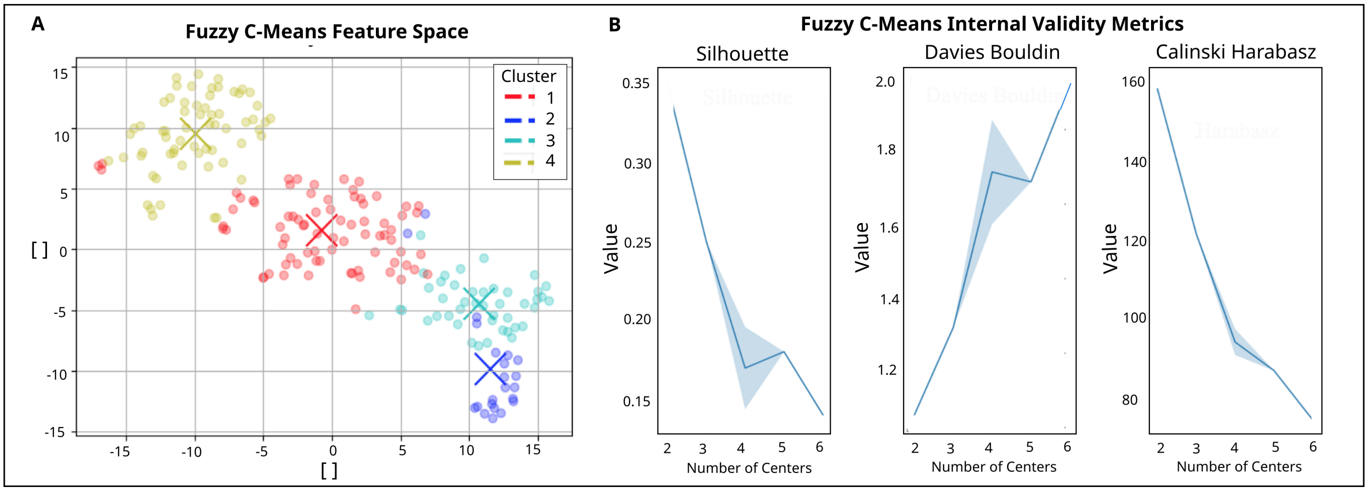


**Supplementary Figure 1. Clustering Results for FCM Analysis**. The left plot (A) shows the feature Space, colourcode represents each cluster. From all data-points, In FCM Clustering, 37.4% lie in Cluster 1, 10.7% in Cluster 2, 21.4% in Cluster 3 and 30.5% in Cluster 4. The right plots (B) show the evolution of the internal validity metrics Silhouette and Davies Bouldin when changing the number of centres. An elbow at k=5 can be seen.

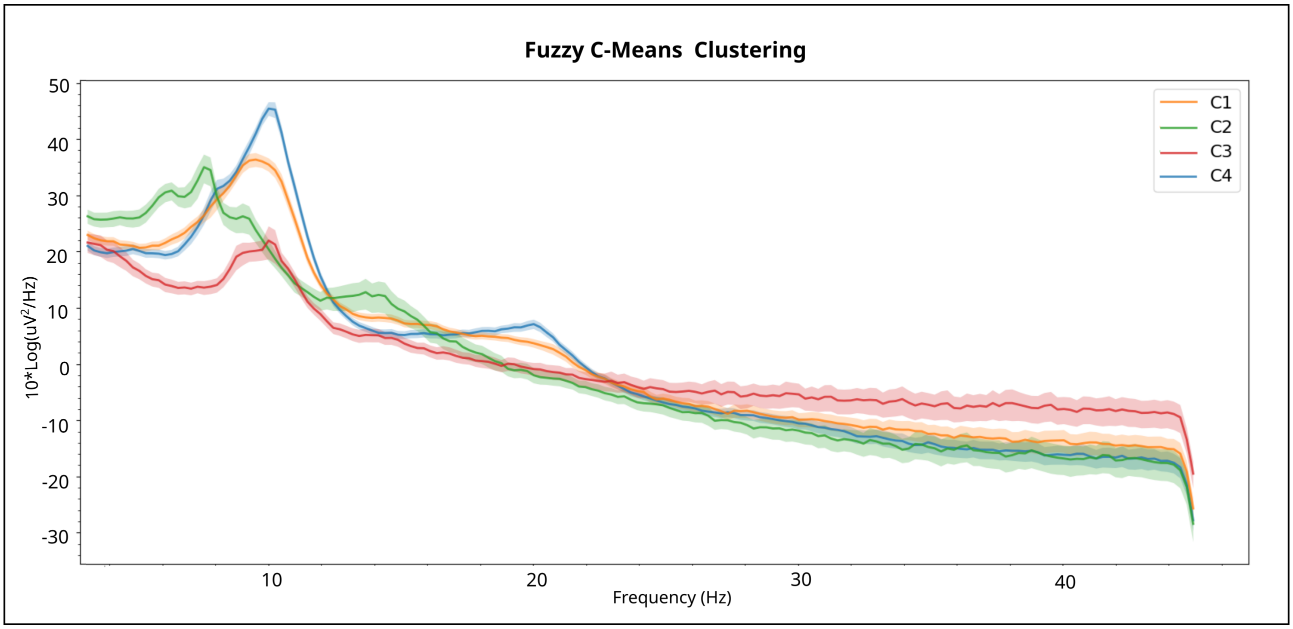


**Supplementary Figure 2. Power Spectrum Plot of each Cluster for FCM Algorithm Results**

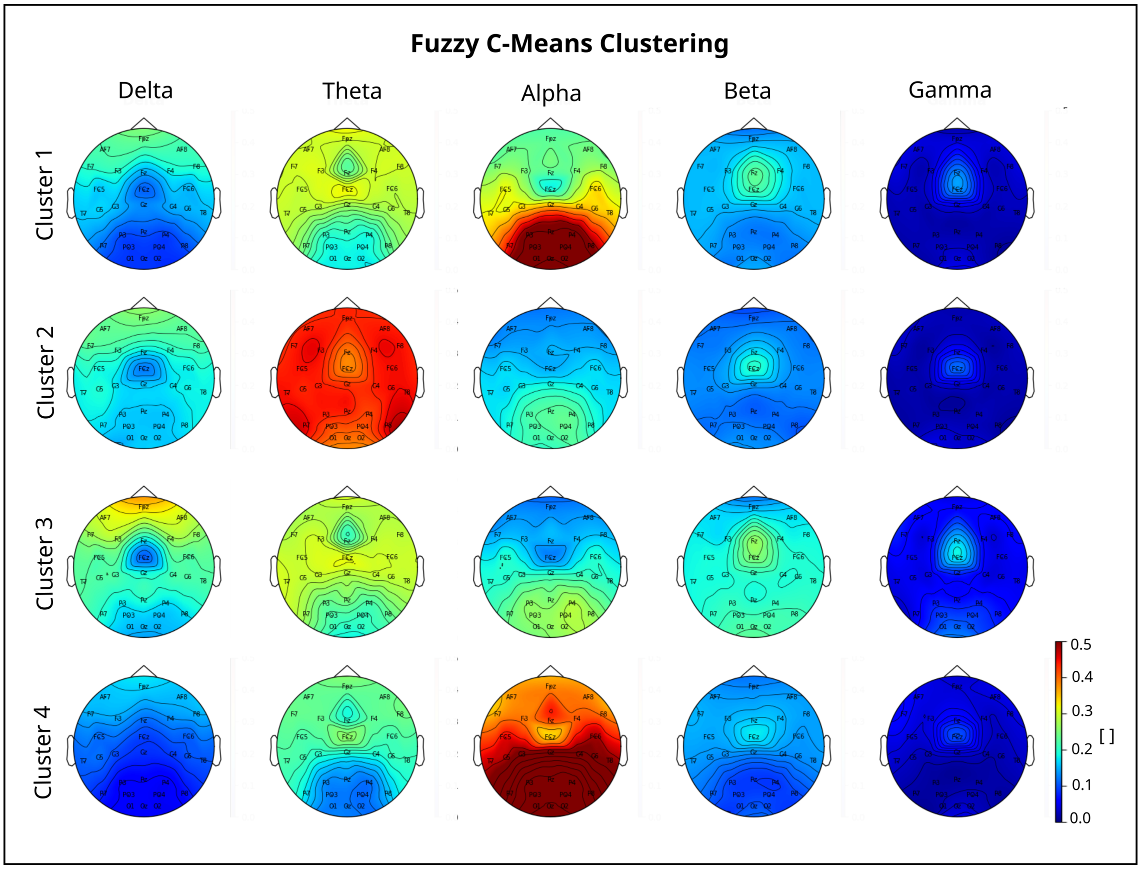


**Supplementary Figure 3.** **Power Spectrum Plot of each Cluster for FCM Algorithm Results**

| Cluster | Group A, Concurrent | Group A, Non-concurrent | Group B, Concurrent | Group B, Non-concurrent |
|---|---|---|---|---|
| 1 | | | | |
| 2 | | | | N-Back Accuracy (+, 0.0287) |
| 3 | | | | Flanker RT (-, 0.0245) |
| 4 | | | | Flanker RT (+, 0.0287) |

**Supplementary Table 1.** **FCM Significant Responders.** These results correspond to significant correlations and significant wilcoxon tests. In parenthesis are: (1) direction of correlation (positive or negative), (2) Wilcoxon p-values. There are significant correlations only in Group B, Non-concurrent, therefore for Wilcoxon Test 5 corrections are done. Stimulation protocol Group B (rIFG), offline mode (non-concurrent task) presents positive response in Clusters 2 and 4 measured with N-Back Accuracy and Flanker RT respectively, and negative response in Cluster 3 assessed with Flanker RT.

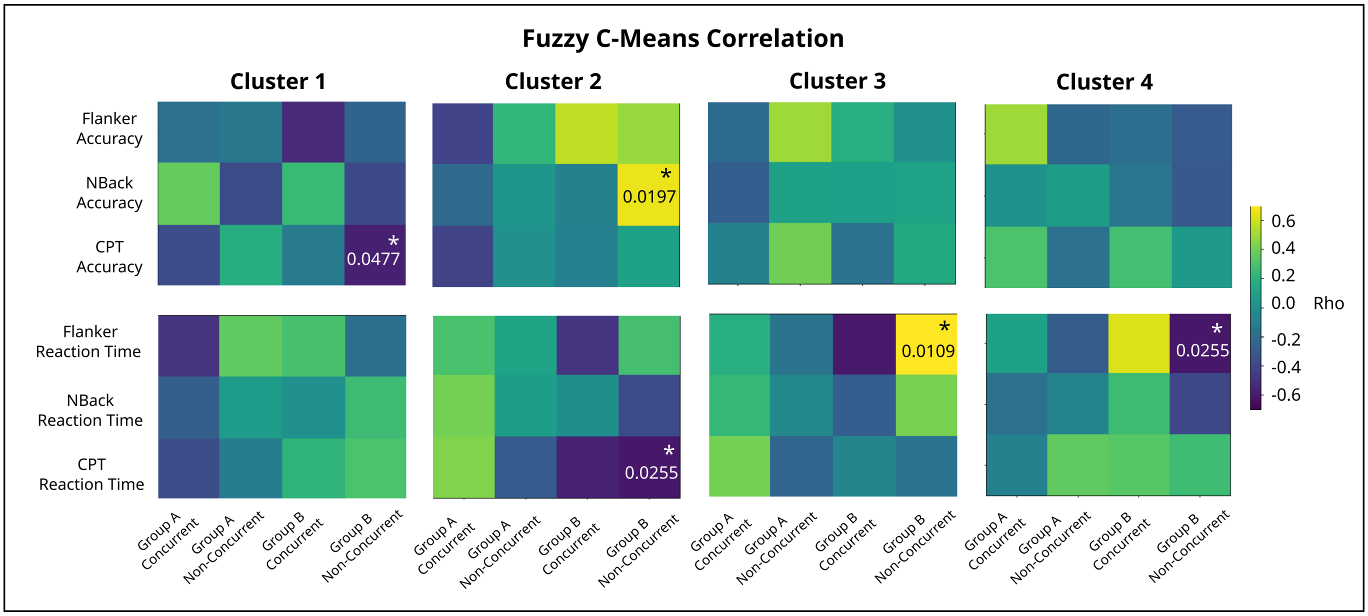


**Supplementary Figure 4.** **Correlation Results for FCM.** Each matrix row corresponds to a specific behavioural task endpoint, and each matrix column corresponds to a particular treatment protocol. The colourbar represents the sign of the correlation (positive or negative) and p-values are written in the significant cells. For Group B, 24 corrections are done; for Group A, 20 corrections are applied.

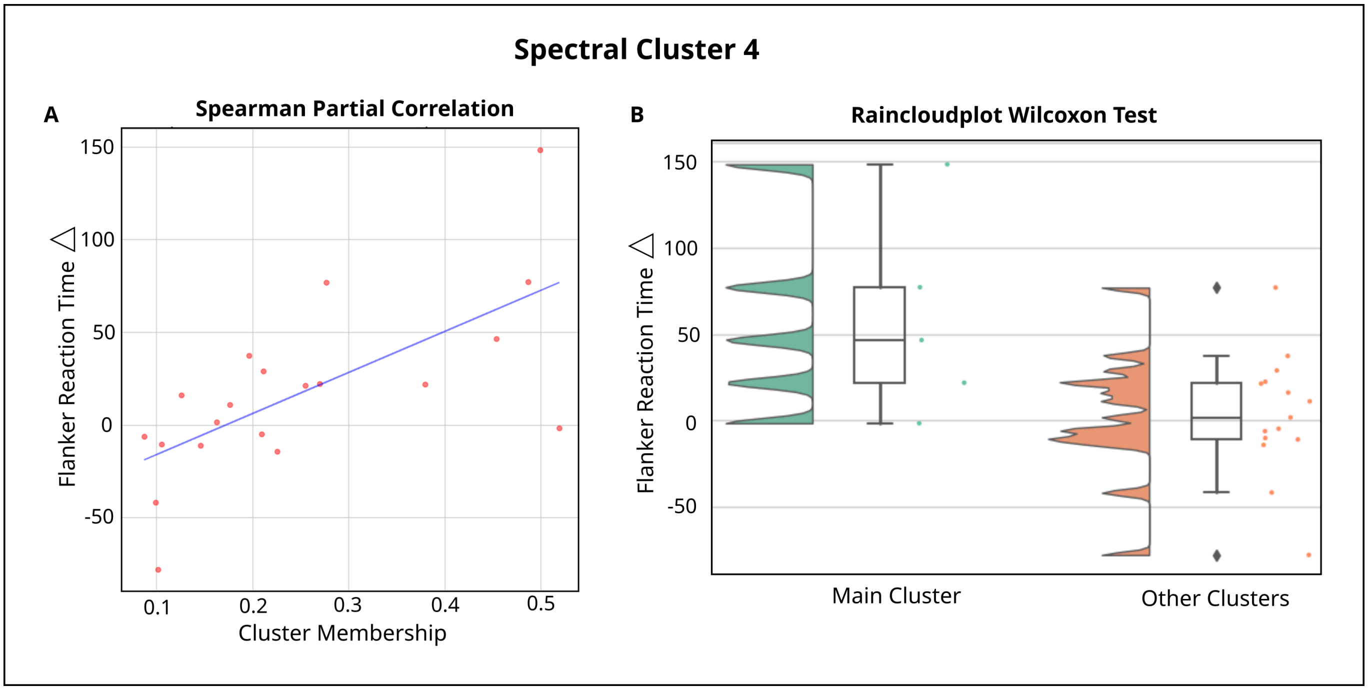


**Supplementary Figure 5.** **Example of Significant responder group for Spectral Clustering (Cluster 3), with rIFG stimulation with no concurrent task, measured with Flanker RT.** (A) Correlation between cluster membership of pre treatment EEG spectral features of all participants and Flanker Reaction Time at the end of the trials. Best-fit line showing least square polynomial fit. (B) Wilcoxon Test between Flanker RT of Cluster 3 Spectral, and the rest of the Clusters. The raw data is visualized as points as well as the probability density, and key summary statistics in the boxplot (median, mean, and confidence intervals).

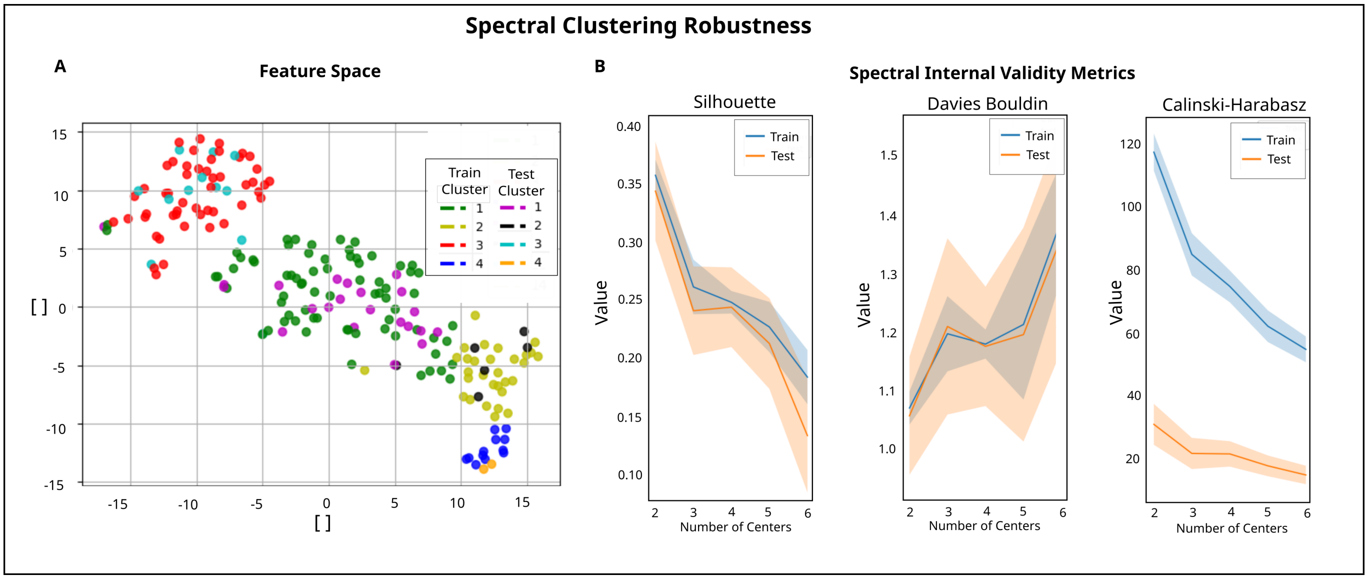


**Supplementary Figure 6. Spectral Clustering Robustness via k-hold-out methodology.** The left plot (A) shows the Feature Space after one iteration, colourcode represents each cluster for each set - train and test. Visual overlap is observed between the clusters assigned to each set. The right plots (B) show the evolution of the internal validity metrics Silhouette and Davies Bouldin when changing the number of centres after 50 iterations. The train and test sets show the same trend.

| Subject ID | Group | Sex | Age |
|---|---|---|---|
| 1f | A | 1 | 15 |
| 2m | A | 2 | 17 |
| 3f | A | 1 | 15 |
| 4m | A | 2 | 14 |
| 5m | A | 2 | 16 |
| 5f | A | 1 | 17 |
| 7m | A | 2 | 13 |
| 7f | A | 1 | 14 |
| 10f | A | 1 | 17 |
| 11f | A | 1 | 10 |
| 12m | A | 2 | 16 |
| 13f | A | 1 | 13 |
| 14m | A | 2 | 15 |
| 15m | A | 2 | 11 |
| 15f | A | 1 | 14 |
| 17f | A | 1 | 14 |
| 17m | A | 2 | 11 |
| 19f | A | 1 | 17 |
| 22m | A | 2 | 13 |
| 25f | A | 1 | 16 |
| 26m | A | 2 | 10 |
| 27m | A | 2 | 12 |
| 28f | A | 1 | 16 |
| 29f | A | 1 | 16 |
| 31f | A | 1 | 15 |
| 33f | A | 1 | 16 |
| 35f | A | 1 | 13 |
| 3m | B | 2 | 17 |
| 4f | B | 1 | 14 |
| 6f | B | 1 | 14 |
| 6m | B | 2 | 15 |
| 8m | B | 2 | 17 |
| 8f | B | 1 | 17 |
| 9f | B | 1 | 13 |
| 10m | B | 2 | 13 |
| 11m | B | 2 | 17 |
| 12f | B | 1 | 12 |
| 13m | B | 2 | 15 |
| 14f | B | 1 | 15 |
| 16m | B | 2 | 11 |
| 16f | B | 1 | 17 |
| 18f | B | 1 | 17 |
| 18m | B | 2 | 10 |
| 19m | B | 2 | 14 |
| 20f | B | 1 | 13 |
| 21m | B | 2 | 11 |
| 22f | B | 1 | 14 |
| 24f | B | 1 | 14 |
| 24m | B | 2 | 11 |
| 25m | B | 2 | 17 |
| 26f | B | 1 | 14 |
| 27f | B | 1 | 14 |
| 28m | B | 2 | 12 |
| 30f | B | 1 | 13 |
| 32f | B | 1 | 14 |
| 34f | B | 1 | 17 |

**Supplementary Table 2.** Clustering participant information. Code per column: sex (1: female, 2: male), task (1: concurrent, 2: non-concurrent), stimulation (1: tDCS, 2: sham), stimulation site (1: lDLPFC, 2: rIFG).

| Subject ID | IQ | Left EHI | Right EHI |
|---|---|---|---|
| 1f | 108 | 1 | 19 |
| 2m | 101 | 1 | 12 |
| 3f | 98 | 3 | 12 |
| 4m | 108 | 2 | 9 |
| 5m | 94 | 2 | 10 |
| 5f | 97 | 1 | 13 |
| 7m | 99 | 19 | 1 |
| 7f | 107 | 10 | 0 |
| 10f | 119 | 1 | 19 |
| 11f | 110 | 20 | 0 |
| 12m | 103 | 2 | 16 |
| 13f | 100 | 0 | 10 |
| 14m | 99 | 0 | 10 |
| 15m | 109 | 3 | 10 |
| 15f | 103 | 0 | 10 |
| 17f | 102 | 0 | 10 |
| 19f | 127 | 0 | 10 |
| 25f | 116 | 12 | 5 |
| 26m | 119 | 0 | 20 |
| 28f | 98 | 1 | 10 |
| 29f | 99 | 4 | 16 |
| 31f | 111 | 5 | 5 |
| 33f | 107 | 0 | 16 |
| 35f | 120 | 12 | 5 |
| 3m | 127 | 3 | 13 |
| 4f | 84 | 0 | 10 |
| 6f | 104 | 0 | 10 |
| 8m | 115 | 3 | 10 |
| 8f | 94 | 2 | 10 |
| 9f | 102 | 0 | 14 |
| 10m | 97 | 0 | 11 |
| 12f | 115 | 1 | 10 |
| 14f | 101 | 10 | 1 |
| 16m | 129 | 4 | 10 |
| 16f | 101 | 0 | 10 |
| 18f | 115 | 4 | 12 |
| 20f | 109 | 10 | 0 |
| 22f | 97 | 1 | 10 |
| 25m | 115 | 0 | 20 |
| 26f | 103 | 1 | 9 |
| 28m | 110 | 2 | 12 |
| 30f | 91 | 3 | 12 |
| 32f | 88 | 2 | 10 |
| 34f | 80 | 0 | 12 |

**Supplementary Table 3.** Correlation Participant information.
